# Mechanisms underlying the efficacy of a rodent model of vertical sleeve gastrectomy — a focus on energy expenditure

**DOI:** 10.1101/2022.03.07.482714

**Authors:** A Stefanidis, CMC Lee, E Greaves, M Montgomery, M Arnold, S Newn, A Budin, CJ Foldi, PR Burton, WA Brown, TA Lutz, MJ Watt, BJ Oldfield

## Abstract

**Background and aims:** Bariatric surgery remains the only effective and durable treatment option for morbid obesity. Vertical Sleeve Gastrectomy (VSG) is currently the most widely performed of these surgeries primarily because of its proven efficacy in generating rapid onset weight loss, improved glucose regulation and reduced mortality compared with other invasive procedures. VSG is associated with reduced appetite, however, the relative importance of energy expenditure to VSG-induced weight loss and changes in glucose regulation, particularly that in brown adipose tissue (BAT), remains unclear. The aim of this study is to investigate the role of BAT thermogenesis in the efficacy of VSG in a rodent model.

**Methods:** Diet-induced obese male Sprague-Dawley rats were either sham-operated, underwent VSG surgery or were pairfed to the food consumed by the VSG group. Rats were also implanted with biotelemetry devices between the interscapular lobes of BAT to assess local changes in BAT temperature as a surrogate measure of thermogenic activity. Metabolic parameters including food intake, body weight and changes in body composition were assessed. To further elucidate the contribution of energy expenditure via BAT thermogenesis to VSG-induced weight loss, a separate cohort of lean rats underwent complete excision of the interscapular BAT (iBAT lipectomy) or chemical denervation using 6-hydroxydopamine (6-OHDA). To localize glucose uptake in specific tissues, an oral glucose tolerance test was combined with an intraperitoneal injection of 2 deoxy-D-glucose (2DG)-14C, administered intraperitoneally. Transneuronal viral tracing was used to identify 1) sensory neurons directed to the stomach or small intestine (H129-RFP) or 2) chains of polysynaptically linked neurons directed to BAT (PRV-GFP) in the *same* animals.

**Results:** Following VSG, there was a rapid reduction in body weight that was associated with reduced food intake, elevated BAT temperature and improved glucose regulation. Rats that underwent VSG had elevated glucose uptake into BAT compared to sham operated animals as well as elevated gene markers related to increased BAT activity (*Ucp1, Dio2, Cpt1b, Cox8b, Ppargc*) and markers of increased browning of white fat (*Ucp1, Dio2, Cited1, Tbx1, Tnfrs9*). Both iBAT lipectomy and 6-OHDA treatment significantly attenuated the impact of VSG on changes in body weight and adiposity in lean animals. In addition, surgical excision of iBAT following VSG significantly reversed VSG-mediated improvements in glucose tolerance, an effect that was independent of circulating insulin levels. Viral tracing studies highlight a patent neural link between the gut and BAT that include groups of premotor BAT-directed neurons in the dorsal raphe and raphe pallidus.

**Conclusion:** Collectively, these data support a role for BAT in mediating the metabolic sequelae, particularly the improvement in glucose regulation following VSG surgery and highlight the need to better understand the contribution from this tissue in human patients.

## Introduction

The efficacy of medicinal treatments for overweight and obesity is improving with the recent introduction of pharmacotherapies drugs and those on the horizon that demonstrate between 5 and 15 % average weight loss^1^. Similarly, lifestyle and behavioral interventions can deliver significant weight loss, however many individuals fail to achieve the levels of weight loss that are required for an effective reduction in comorbidities, which is often only associated with long term compliance^2^. By comparison, bariatric surgery remains the only effective and durable treatment for morbid obesity providing up to 30% body weight loss^3, 4^. Vertical sleeve gastrectomy (VSG) is currently the most widely performed weight loss surgery based on its utility in producing reliable weight loss, improved glucose regulation and only moderate levels of surgical complications and reoperation^5^. The procedure involves a longitudinal resection of 80% of the greater curvature of the stomach to generate a tubular duct or “sleeve” with preservation of the intestinal anatomy.

Reduction in body weight following VSG has been attributed to enhanced gastric emptying^6^, accelerated intestinal transit^7^, and sustained changes in gut hormones that promote satiety^8, 9^. A reduced concentration of the orexigenic hormone ghrelin after VSG has consistently been reported^10^, an effect that has been attributed to the resection of much of the gastric fundus where ghrelin is produced^11^. Furthermore, elevations in postprandial levels of anorexigenic hormones glucagon like peptide-1 (GLP-1) and peptide YY (PYY) following VSG have been demonstrated in both animal^12^ and human studies^8, 13^. It is possible that these hormones act centrally to cause satiety but it is also likely that they act locally in a paracrine fashion, along with other gut related stimuli including shifts in gut microbiota and bile acids, on vagal afferent neurons terminating in the stomach and duodenum^14, 15^.

The other major metabolic outcome of VSG, the immediate postsurgical improvement in glycemic control has, in common with the presumptive drivers of reduced food intake and body weight, been proposed to be mediated by elevated levels of postprandial GLP-1 and PYY^13^. Other potential underlying mechanisms involve surgically-induced reductions in the elusive foregut “anti-incretins” ultimately improving glucose regulation^9^, incompletely defined intestinal adaptations to surgery^16^ and complex interactions between bile acids, fibroblast growth factor 19 (FGF-19) and the gut microbiome^17^.

In addition to the potential mechanisms listed above that may underlie the efficacy of VSG, the recruitment of brown adipose tissue (BAT) and “browned” white adipose tissue has been implicated, particularly in relation to post bariatric surgical *weight loss*^18^. These data, derived from both rodent studies and clinical interventions show an involvement of BAT and beige adipose tissue (BeAT) in mediating the efficacy of bariatric surgery with a consistent trend toward greater recruitment of both classical BAT and BeAT after VSG than after Roux-en-Y, with the latter promoting changes predominantly in BeAT in visceral and subcutaneous depots. Such studies, while intriguing, have focused on weight loss rather than glucose regulation^19, 20^. Given the well-established and remarkable contribution of brown and beige fat to the sequestration of triglycerides from the blood it is surprising that, with notable exceptions in both human^21^ and rodent^22, 23^ studies of VSG, the potential for elevated activity of BAT and BeAT has not been highlighted as an explanation for the dramatic improvement in glucose regulation immediately after some resectional bariatric surgeries. In this respect, it has been shown in patients that there an increase in the fractional uptake of the fatty acid radioisotopes 18F-FTHA and non-esterified fatty acids (NEFA) in the supraclavicular BAT after VSG and RYGB^21^ and in rodent studies there is increased uptake of the glucose analogue ^18^F-FDG into BeAT after VSG in chow-fed mice.

The present study, conducted in a rat model of VSG, aims to elucidate the importance of brown and beige fat in VSG-mediated metabolic benefit, both in terms of body weight and glycemic control. Importantly, this data highlights a potential vagal–brain–BAT pathway that could provide a neural substrate for such a reflex action and provides insights into the means by which BAT/BeAT is recruited and effective in contributing to improved metabolic outcomes after VSG.

## Methods

### Animals

Male Sprague-Dawley rats (*n*=170; 4-12 weeks old; Monash Animal Services, Australia) were housed under a 12h: 12h light/dark cycle and temperature-controlled environment (27±1°C). Rats had *ad libitum* access to food and water during the experimental period unless specified. Rats were either made obese (diet-induced obese, DIO) by placing them on a on a high fat diet (45% kcal from lipids, 23.5% total fat, SF04–001; Specialty Feeds, WA, Australia) for 14-20 weeks or maintained on a standard chow diet (12% kcal from lipids, 4.6% total fat; Rat and Mouse cubes, Specialty Feeds, WA, Australia). DIO rats continued to have *ad libitum* access to water and high fat diet following surgery (VSG or sham) unless specified. All experimental procedures in this study were reviewed and approved by the Monash Animal Resource Platform animal ethics committee (MARP-1 AEC;17793) and animals were treated in accordance with the *Australian Code of Practice for the Care and Use of Animals for Scientific Purposes.*

#### Characterization of metabolic outcomes following VSG

Metabolic parameters were assessed in DIO rats (n=20) that had undergone sham-operation, VSG surgery or a pair-feeding paradigm over a 40 day monitoring period including the assessment of daily changes in body weight and food intake (up to day 31), changes in body composition (pre-vs post-surgery using dual energy x-ray absorptiometry, DEXA, Discovery DXA system, Hologic, NSW, Australia) and glucose regulation (oral glucose tolerance rest, oGTT, 1.5g/kg lean mass).

#### Changes in energy expenditure following VSG

A separate cohort of DIO rats (n=15) that had undergone sham-operation, VSG surgery or the pair-feeding paradigm were assessed for changes in BAT temperature 10-14 days following surgery, via an implanted thermistor tip (TA10TA-F40, Data Sciences, USA) positioned between the two interscapular lobes of BAT, as a marker of thermogenic activity in BAT. Other elements of energy expenditure (n=16) including oxygen consumption, energy expenditure substrate utilisation (respiratory exchange ratio; RER) were also assessed using an indirect calorimetry system (LabMaster; TSE-systems, Bad Homburg, Germany) 30-35 days after VSG or sham surgery. Changes in UCP1 expression (n=12), as a marker of thermogenic activity was assessed using Western blot analysis ^24^.

#### Changes in glucose regulation following VSG

In order to assess the impact of VSG on glucose homeostasis, an oGTT was combined with an intraperitoneal injection of [1-^14^C]-deoxy-d-glucose tracer (40μCi) performed 35-40 days following surgery to determine tissue specific glucose uptake (n=26) in sham- and VSG-operated DIO rats and compared to age-matched chow-fed rats. BAT and iWAT samples (n=20) were also assessed for changes in the gene expression of markers of increased brown fat activity and the browning of WAT using quantitative RT-PCR^24^.

#### Impact of disrupting BAT on VSG-induced weight loss

Chow-fed rats (n=32) were either sham-operated or underwent VSG surgery. Rats were then subdivided into six groups, either serving as controls with intact and functional BAT (Sham+Intact and VSG+Intact), receiving complete excision of the interscapular BAT (Sham+iBAT lipectomy; VSG+iBAT lipectomy) or bilateral injections of 6-hydroxydopamine (6-OHDA) into the interscapular BAT (Sham+6-OHDA; VSG+6-OHDA). Changes in body weight, food intake and composition were monitored over the subsequent 14 days.

#### Impact of disrupting BAT on VSG-induced changes in glucose regulation

In order to assess the importance of iBAT in the changes in glucose tolerance following VSG, a separate cohort of DIO rats (n=10) had oGTTs (1.5g/kg lean mass) performed six weeks following sham or VSG surgery and compared to chow-fed controls (n=6). Rats then underwent complete excision of their iBAT and oGTTs were repeated at least 7 days following surgical recovery.

#### Changes in neural activity following the infusion of nutrient and non-nutrient loads

To elucidate whether the effect of VSG on neural activation was in response to volume or nutrient content, a cohort of chow-fed rats (n=24) were either sham-operated or underwent VSG surgery and within these groups, rats were further divided into three groups receiving different stimuli. Following a six-hour fast, rats were either 1) handled only, 2) received an intragastric infusion of 2 mL water, or 3) received 2mL of a mixed nutrient meal (4kcal, Ensure, Abbott Nutrition, NSW, Australia). Two hours later, rats were anaesthetized and perfused transcardially with physiological saline 4% paraformaldehyde and processed using standard immunohistochemical approaches to identify Fos protein expression ^25^.

#### Identification of neuroanatomical pathways directed from the stomach to the brain

A recombinant of the alpha herpesviruses, H129 that expresses red fluorescent protein (RFP;H129-RFP) was used to conduct neuroanatomical tracing of VANs and their neural connections immediately after sham (n=5) or VSG (n=5) surgery. Animals were allowed to survive for up to five days post-inoculation to ensure early arrival of the virus at the region at which point they were perfused transcardially and nodose ganglia and brains collected.

#### Identification of synaptic relays between sensory pathways from the stomach or small intestine and efferent outflows to the BAT

A combination viral approach was adopted that involved the injection of H129-RFP into the stomach or duodenum (small intestine) and PRV-GFP into the iBAT in the same animal. Injections into the stomach (n=8) or duodenum (n=8) were first performed (as above) followed by BAT injections which were staggered by a day to ensure simultaneous arrival of both viruses within discrete brain regions. Following survival of up to five days, rats were perfused transcardially and brains collected.

Additional experimental details are provided in the Supplementary Methods and include extended information related to animal preparation and surgery, experimental protocols and reagents used.

### Statistical analysis

Statistical analyses were conducted using Prism (GraphPad Software (v9.0), Inc., San Diego, CA). Comparative statistical analyses were performed using a repeated measures analysis of variance (ANOVA) for the assessment of body weight, food intake, BAT temperature, glucose tolerance and neural activity (Fos immunoreactivity). Bonferroni *post-hoc* tests were performed to determine differences between individual treatment groups. *Post-hoc* analysis was performed when a treatment/surgery by time interaction was observed. Differences in protein and gene expression were assessed using a two-tailed unpaired T-test. In the case of assessment of the number of Fos-positive neurons, a repeated measures ANOVA was used to determine changes throughout the NTS. When a treatment/surgery interaction was observed, a Bonferroni *post-hoc* test was performed to assess the level where differences exist. Values of *p* < .05 were regarded as statistically significant. All results are presented as mean ± SEM.

## Results

### Effect of VSG on energy balance in DIO rats

#### Body weight

Sham-operated rats showed a transient decrease in body weight over the first 5 days post-operatively, which was gradually regained. Over the first 10 days, rats that underwent VSG had the most pronounced reduction in body weight compared to sham-operated controls whereas pairfeeding caused a shift in the rate of body weight loss over this initial period (*F*_(20,160)_=15.33, *p*_surgeryXtime_<.0001; Fig.1B). Compared to the sham-operated controls, the reduction in body weight in VSG-operated and pairfed rats persisted (*F*_(2,16)_=16.85, *p*_surgery_=.0001, *F*_(60,480)_= 12.23, *p*_surgeryXtime_<.0001;Fig.1C) over the remainder of the measurement period.

**Figure 1.**
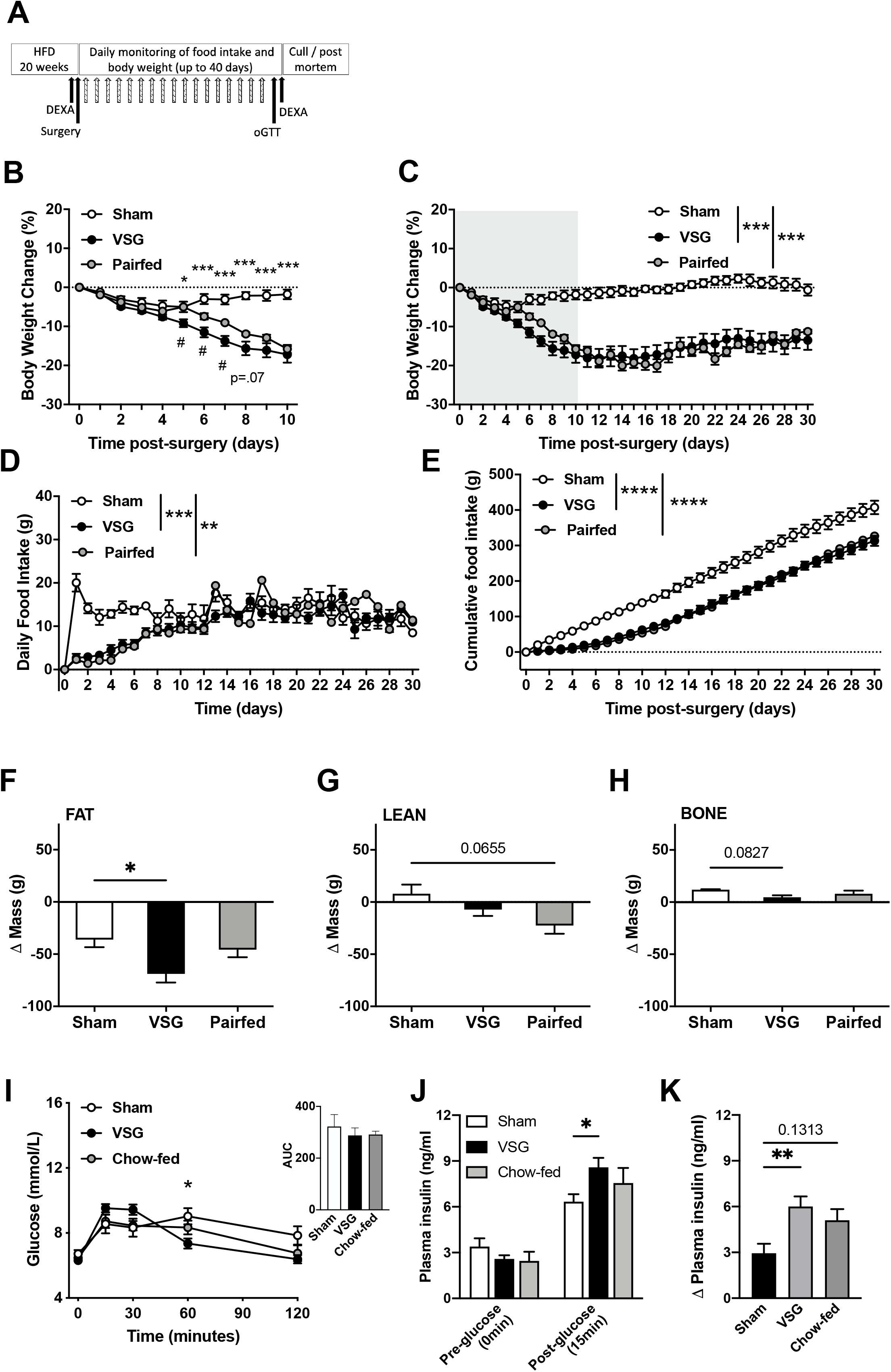
Changes in body weight, food intake, body composition and glucose tolerance following VSG in DIO rats. (A) Study timeline, (B) and (C) comparison of the effects of sham operation (n=5), VSG (n=9) and pair-feeding (n=5) (A) on body weight change (%) over the rapid weight loss phase (A) and over the 31-day monitoring period (B), (D) daily food intake (g), (E) cumulative food intake (g) in DIO rats. Changes in body composition including (F) fat mass (g), (G) lean mass (g) and (H) bone mass (g) relative to baseline (pre-surgical) levels. Blood glucose levels and AUC analysis (I) following glucose (1.5g/kg lean mass) in sham(n=8) or VSG(n=9) operated rats compared to chow-fed(n=6) controls. (J) Plasma insulin (ng/ml) at baseline and 15minutes following glucose and (K) change in insulin levels relative to baseline concentration. n=4–9. *P<.05,**P<.01, ***P<.001,****P<.0001,* in B, I, comparing sham vs VSG, #in B,VSG vs pairfed.

#### Food intake

Food intake was significantly reduced VSG surgery compared to sham-operated controls regardless of being expressed as either average daily (*F*_(2,16)_=11.08, *p*_surgery_=.001,*F*_(60,480)_=5.216, *p*_surgeryXtime_<.0001, Sham vs VSG, *p*<.001; Fig.1D) or cumulative food intake (*F*_(2,16)_=24.24 *p*_surgery_=.001,*F*_(60,480)_=6.262, *p*_surgeryXtime_<.0001, Sham vs VSG, *p*<.0001; Fig.1E). This reduction was most pronounced in the first 7 days post-VSG surgery and was also evident between pairfed and sham groups (Sham vs Pairfed, *p*<.01; Fig.1D, E). Interestingly, the food intake was similar between sham and VSG-operated rats over the second half of the monitoring period highlighting possible alternate mechanisms in the sustained weight loss induced by VSG.

#### Body composition

DEXA scans revealed a significant reduction in fat mass (*F*_(2,15)_=*4.445*, *p*_surgery_<.0001) in rats that underwent VSG as compared to sham controls (Sham vs VSG, p<.05) (Fig. 1F). Pair-feeding tended to preferentially reduce lean mass (*p*=.0655) whereas a tendency for a loss in bone mass was apparent following VSG (*p*=.0827) but not pair-feeding (Fig.1F–H).

#### Glucose tolerance testing

In response to an oral glucose load (1.5g/kg lean mass), there was shift in glucose regulation where VSG initially enhanced blood glucose levels following VSG compared to sham operated controls (15 and 30min, p<.05) (*F*_(8,80)_=4.126, *p*_surgeryXtime_<.001) followed by a substantial decline in blood glucose levels in the VSG group compared to sham-operated controls at the 60 min timepoint, where blood glucose levels were significantly higher in the sham operated group and did not return to baseline levels (Figure 1H). Interestingly, VSG did not impact baseline insulin concentrations (pre-glucose), however, as expected plasma insulin concentrations were higher for the VSG-operated rats compared to sham-operated controls 15 min following the glucose gavage (*F*_(2,20)_=5.955, *p*_surgeryXtime_<.001, Sham vs VSG, *p*<.05). Furthermore, the extent to which glucose is capable of increasing plasma insulin concentrations relative to baseline levels was significantly enhanced in the VSG group (*F*_(2,20)_=5.955, *p*_surgery_<.01,Sham vs VSG, *p*<.01) and interestingly, the VSG mediated response was indistinguishable from the chow-fed group (VSG vs chow-fed, *p* >.9999) (Fig.1J, K).

### The role of energy expenditure on VSG-induced weight loss

Similar to above, VSG resulted in a significant reduction in body weight gain compared to sham operated controls while the rate of weight loss tended to be attenuated in the pairfed group compared to that caused by VSG (*F*_(28,168)_=9.052, *p*_surgeryXtime_ <.0001, *F*_(2,12)_=19.50, *p*_surgery_<.001)(Fig.2A).

**Figure 2.**
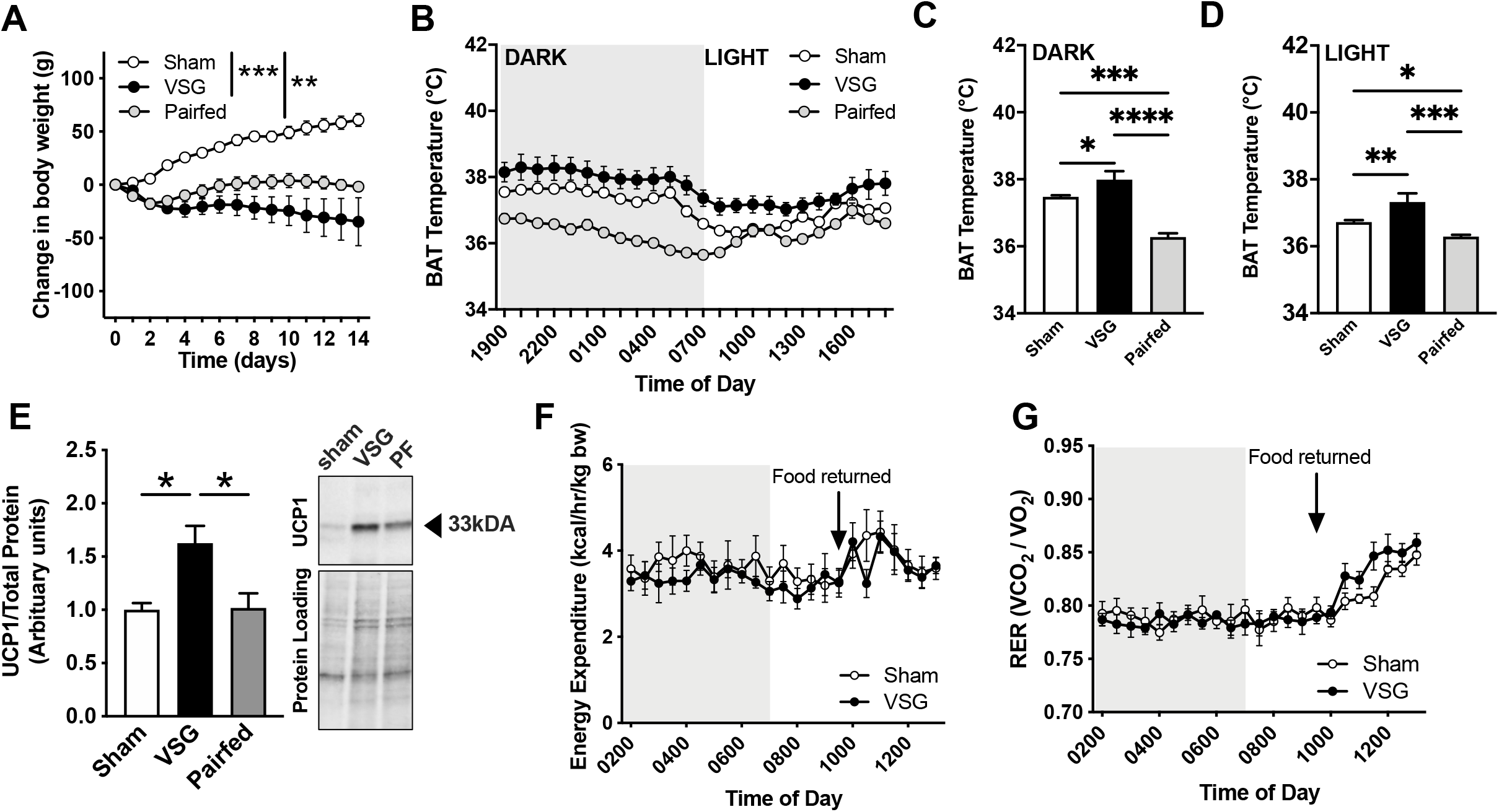
Changes in energy expenditure following VSG in DIO rats. Changes in (A) body weight (g) and (B) mean hourly BAT temperature of DIO rats during the (C) dark and (D) light period following sham operation (n=5), VSG (n=5) or pair-feeding (n=5). (E) Western blot detection of UCP1 protein expression in BAT and a representative blot is shown (n=6-9/group). Changes in (F) Energy expenditure and (G) RER during a fast and upon refeeding (0930h) (n=8/group). n=5–8. *P<.05, **P<.01, ***P<.001, ****P<.0001.

#### BAT thermogenesis

BAT temperature exhibited a circadian rhythm such that it was elevated during the dark and reduced during the light phases (Fig. 2B). Rats that underwent VSG showed a significant elevation in BAT temperature relative to sham controls (*F*_(48,276)_=7.335, *p*_surgeryXtime_<.0001, *F*_(2,12)_=15.76, *p*_Surgery_<.0001) in both the dark (*F*_(2,12)_= 28.30, *p*_surgery_<.0001, Sham vs VSG *p*=.018,Fig.2C) and light periods (*F*_(2,12)_=11.08, *p*_surgery_=.0019, Sham vs VSG *p*=.0504, Fig.2D). In contrast, pairfed animals had a marked reduction in BAT temperature compared to sham-operated rats (*p*=.0002, Fig.2B) during the dark (*p*<.05, Fig.2C) and light periods (*p*<.05,Fig.2D), highlighting an effect on BAT thermogenesis that extended beyond food intake. Furthermore, western blot analysis demonstrated a significant increase in UCP1 expression in BAT in VSG-operated rats compared to both sham controls and pairfed rats (*p*<.05, Fig.2E).

Basal oxygen consumption, energy expenditure and RER were unaffected by VSG surgery when compared to sham surgery (data not shown). Similarly, energy expenditure (*F*_(22,308)_=.7397, *p*_surgeryXtime_=.7973, *F*_(1,14)_=.4534, *p*_surgery_=.5117,Fig.2F) and RER (*F*_(22,308)_=1.499, *p*_surgeryXtime_=.072, *F*_(1,14)_=.1755, *p*_surgery_=.6816,Fig.2G) were unaffected by fasting however, when assessed over the three hours following the return of food, RER was elevated (*F*_(6,84)_=1.617, *p*_surgeryXtime_=.1524, *F*_(1,14)_= 4.704, *p*_surgery_<.05) in the rats that have undergone VSG surgery, suggesting a preferential shift towards carbohydrate rather than fat oxidation (Fig.2G).

### Changes in tissue specific glucose uptake

Thirty minutes following the injection of the radiotracer, there was a striking increase in ^14^C-2DG content in BAT (*F*_(2,15)_=5.616,p_surgery_=.0151; Sham vs VSG, p=.0184,Fig.3A) and tended to increase in rWAT (*F*_(2,16)_=2.664, *p*_Surgery_=.1003, Fig.3D) but not iWAT or eWAT (Fig.3B,C) following VSG surgery when compared to those undergoing sham surgery. Importantly, the glucose uptake in these tissues was indistinguishable from chow-fed control rats (Fig.3A,D). Accumulation of tracer in skeletal muscle (gastrocnemius and soleus), heart and kidney were similar between sham and VSG-operated rats, and VSG did not restore high fat diet induced impairments in glucose uptake in muscle or heart (Fig.3E-H).

**Figure 3.**
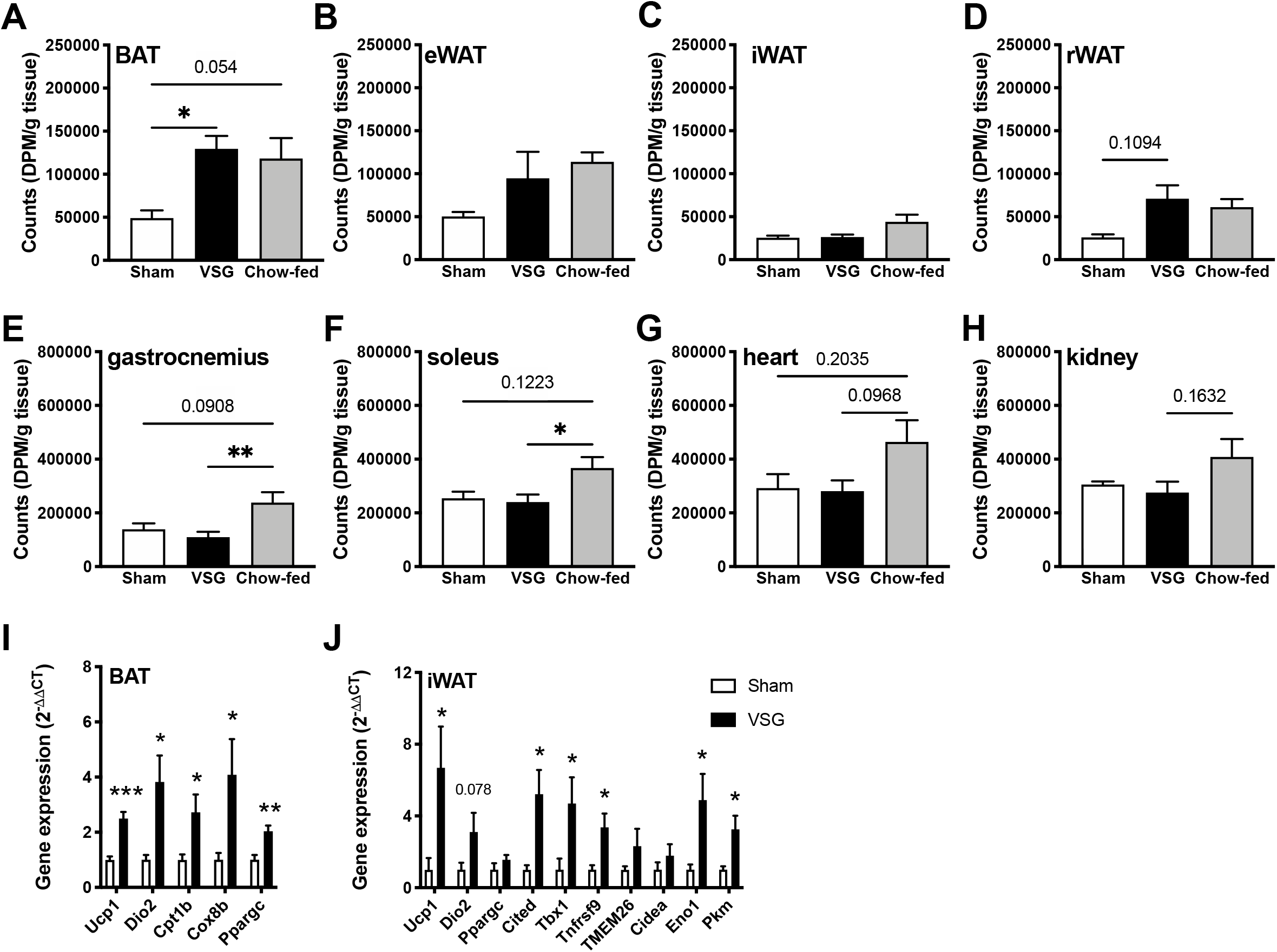
Changes in glucose-stimulated glucose uptake following VSG and gene expression of brown and beige cell markers. Glucose uptake 30 minutes following the administration of glucose (1.5g/kg lean mass) in (A) BAT, (B) eWAT, (C) iWAT, (D) rWAT, (E) gastrocnemius and (F) soleus skeletal muscle, (G) heart and (H) liver following sham (n=8) or VSG (n=9) surgery compared to chow-fed (n=6) controls. Gene expression of brown and beige cell markers in (I) BAT and (J) iWAT (n=5–9). *P<.05, **P<.01, ***P<.001.*in I and J, compared to Sham. Abbreviations: BAT, brown adipose tissue; eWAT, epidydminal white adipose tissue; iWAT, inguinal white adipose tissue; rWAT, retroperitoneal adipose tissue.

### Changes in the expression of genes involved in the regulation of BAT activity and the browning of white adipose tissue

Gene expression analysis showed an elevation in markers related to increased BAT activity including *Ucp1* (*t*_(15)_=5.403, *p*<.0001), *Dio2*(*t*_(14)_=2.556, *p*<.05), *Cpt1b*(*t*_(15)_=2.432, *p*<.05), *Cox8b*(*t*_(15)_=2.207, *p*<.05), and *Ppargc*(*t*_(15)_=3.767, *p*<.005), in BAT in rats following VSG compared to sham operated controls (Fig.3I). Similarly, in iWAT, markers of BAT activity *Ucp1* (*t*_(10)_=2.376, *p*<.05) and *Dio2*(*t*_(9)_=1.990, *p*=.0778) and genes involved in the browning of WAT were significantly elevated [*Cited1*(*t*_(10)_=3.065, *p*=.0119), *Tbx1* (*t*_(10)_=2.317, *p*<.05), *Tnfs9*(*t*_(10)_=2.887, *p*=.0162)] or tended to increase (*TMEM26, Cidea*) following VSG compared to sham operated controls. In addition, the expression of *Eno1*(*t*_(9)_=2.873, *p*=.0184) and *Pkm2*(*t*_(10)_=3.134, *p*=.0120), markers of glycolytic beige cells, were significantly elevated in iWAT following VSG compared to sham surgery (Fig.3J).

### Metabolic consequences of surgical excision and chemical denervation of iBAT

#### Body weight and composition

Changes in body weight were assessed over the 13-day treatment period following either complete excision of the iBAT or denervation using 6-OHDA. There was a significant reduction in body weight in rats that underwent VSG surgery regardless of whether BAT was intact, excised or chemically denervated compared to sham controls. However, the extent of weight loss following VSG relative to sham controls (*F*_(13,104)_=14.53,psurgeryXtime<.0001, *F*_(1,8)_=31.84,p_surgery_=.0005, Fig.4A) was reduced by approximately 50% after both complete excision of the iBAT (*F*_(13,117)_=5.401,p_surgeryXtime_<.0001,*F*_(1,8)_=9.955,p_surgery_=.0116,Fig.4B) or 6-OHDA treatment causing chemical denervation (*F*_(13,130)_=7.964,p_surgeryXtime_<.0001,*F*_(1,10)_=9.261,p_surgery_=.0124,Fig.4C). VSG-induced weight loss in animals with undisrupted BAT was associated with a significant reduction in fat mass relative to sham controls when expressed as a change in the total percentage body fat (*p*<.05, Fig.4E). These effects were absent in VSG animals with surgically and chemically denervated BAT relative to their controls (Fig.4F,G).

**Figure 4.**
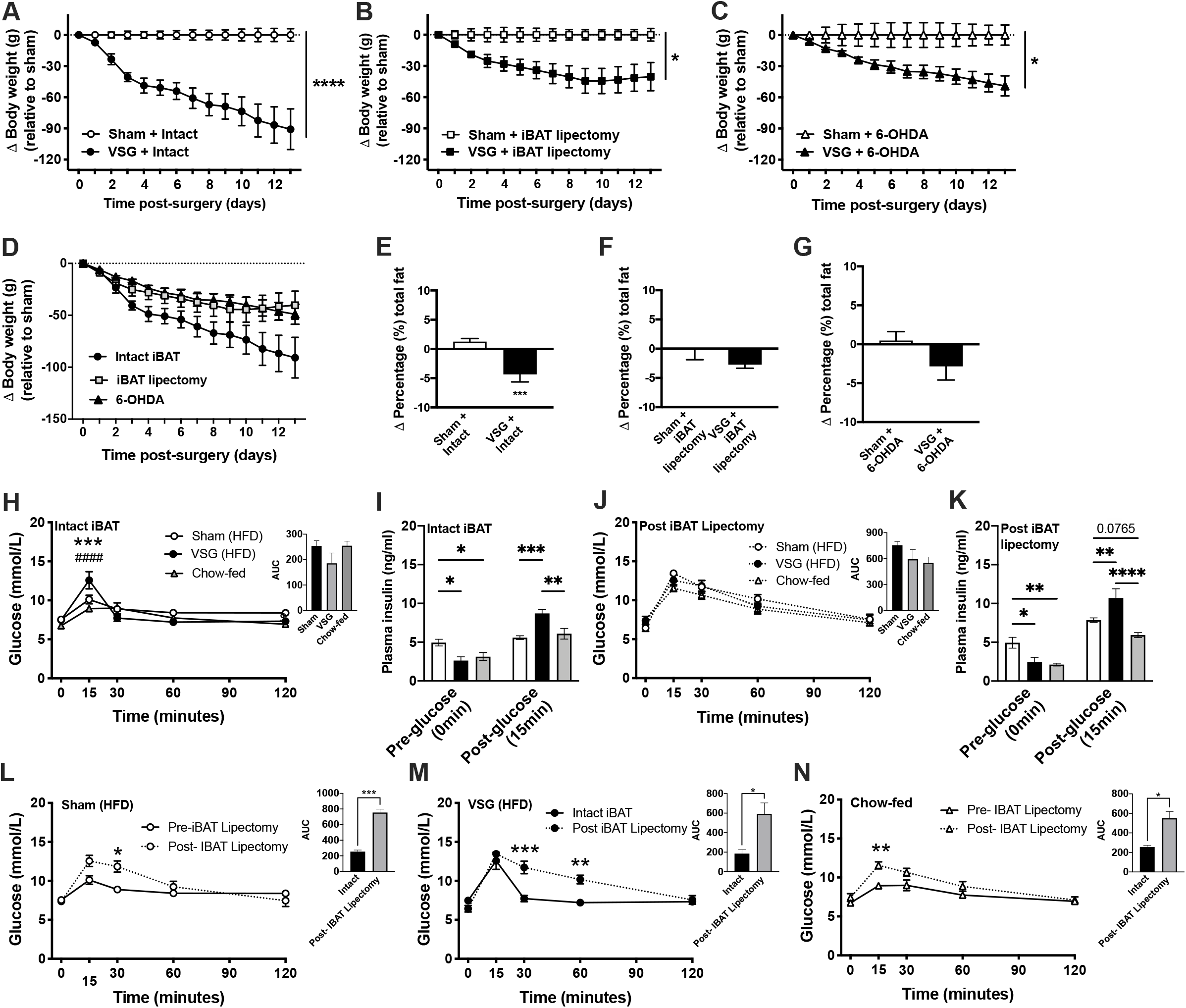
Changes in body weight (g) relative to the respective sham operations in lean rats over 13 days post-surgery. Body weight was monitored following (A) sham operation (n=5) or VSG (n=5) with intact iBAT function, (B) complete excision of iBAT [sham+iBAT lipectomy(n=5), VSG+iBAT lipectomy (n=5)] and (C) chemical denervation of iBAT using 6-OHDA [(sham+ 6-OHDA (n=5), VSG+6-OHDA (n=7)]. Changes in the proportion of fat (%) in rats with (E) intact iBAT function, (F) complete excision of iBAT and (G) chemical denervation of iBAT. Blood glucose (mmol/L) during oGTT with AUC in DIO rats following sham (n=6) or VSG (n=4) surgery and chow-fed controls (n=6) with (H) intact iBAT and (J) following iBAT excision. Plasma insulin levels during oGTT at baseline (pre-glucose) and 15 minutes following oral glucose in rats with (J) intact iBAT and (K) following surgical excision of the iBAT. Comparison of (L) sham-operated, (M) V SG-operated and (N) chow-fed rats prior to and following iBAT excision. *P<.05,**P<.01,***P<.001,****P<.0001.* in H, Sham vs VSG;# in H, VSG vs Chow-fed.

#### Glucose regulation

In order to assess the role of iBAT in the changes in glucose homeostasis that proceed VSG, oGTTs were performed in DIO rats following VSG, before and after excision of the iBAT. In response to the glucose load, there was a peak in blood glucose levels that occurred at 15 mins in rats with and without iBAT (Fig. 4H,4J). Following VSG, there was the characteristic elevation in blood glucose levels following the glucose load at 15 minutes, which was rapidly cleared to baseline levels over the subsequent 15 minutes (*F*_(8,52)_=7.329,p_surgeryXtime_<.0001, F_(2,13)_=2.776,p_surgery_=.0991;Fig.4H). By comparison, following iBAT lipectomy, there was no detectable differences in glucose tolerance between sham, VSG-operated and chow-fed rats (*F*_(8,52)_=1.217, *p*_surgeryXtime_=.3075, *F*_(2,13)_=1.025 *p*_surgery_=.3860; Fig.4J). Interestingly, the reduction in fasting insulin levels and increased capacity for a glucose load to increase plasma insulin concentrations in VSG operated rats was similar in rats with (*F*_(2,11)_=37.19, *p*_surgeryXtime_ <.01,Fig.4J) and without iBAT (*F*_(2, 11)_=16.73, *p*_surgeryXtime_=.0005, Fig.4L). When oGTTs were compared in each of the groups prior to and following iBAT lipectomy, there was a marked reduction in the clearance of blood glucose, an effect that was also apparent in VSG-operated rats at the 30 and 60 min timepoints (*F*_(4,12)_=8.002, *p*_iBATlipectonyXtime_<.005, *F*_(1,3)_ = 5.367, *p*_iBATlipectomy_=. 1034; Fig.4M).

### Changes in neural activity following VSG

#### Acute infusion of either nutrient or non-nutrient loads into the VSG stomach

While intragastric infusion of water had no effect on Fos protein expression at any level of the NTS in sham operated rats, the impact of pressure arising from water infused into the VSG stomach produced a significant elevation in the numbers of Fos-labelled neurons in the NTS (Fig.5), primarily in the mid (*F*_(2,14)_=41.52, *p*_surgeryXinfusion_<.0001, *F*_(1,14)_=117.8, *p*_infusion_<.0001, *F*_(2,14)_=54.10, p_surgery_<.0001; Fig.5B) and caudal levels (*F*_(2,14)_=9.544,p_surgeryXinfusion_<.005, *F*_(1,14)_=56.64,p_infusion_<.0001, *F*_(2,14)_=28.00,p_surgery_<.0001;Fig.5C). By comparison, infusion of the same volume of mixed-meal (Ensure) resulted in significantly increased numbers of Fos-expressing neurons at all levels of the NTS in VSG rats compared to those that were handled only (Fig.5A-D). The number of Fos-positive neurons in the rostral and mid portions of the NTS after Ensure infusion was approximately twice that detected after the same volume of water (Fig.5A,B), an effect that was similar when numbers of activated neurons were considered across the nucleus as a whole (Fig.5D). These data are consistent with a possible effect of volume and nutrients to the recruitment of neurons in the NTS after infusions into the VSG stomach.

**Figure 5.**
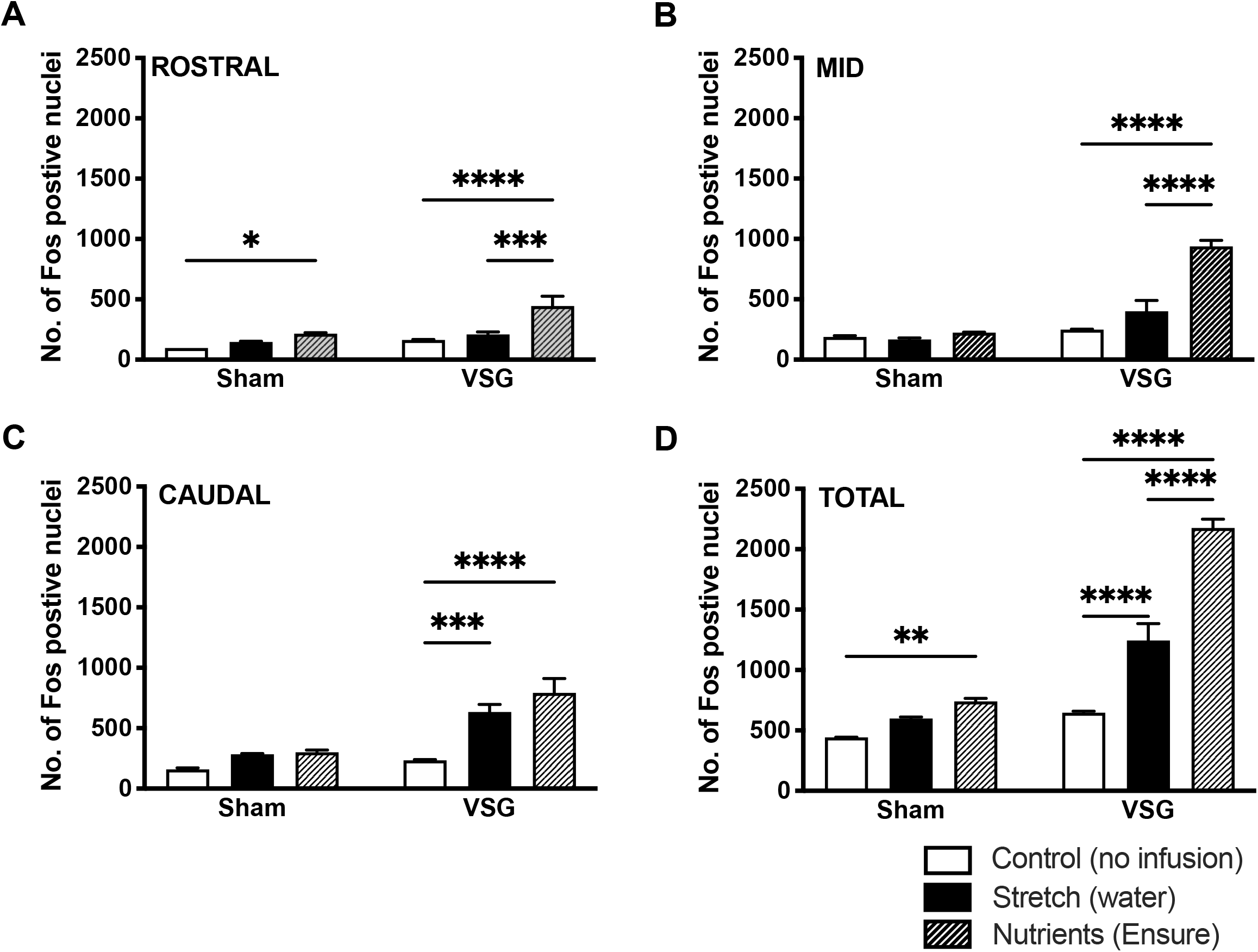
Quantitative assessment of the extent of labelling of neurons containing Fos in chow-fed rats following sham surgery + Control (no infusion), sham surgery + stretch (water), sham + nutrients (Ensure) or VSG + Control, VSG + Stretch, VSG + Nutrients. Counts were made through the (A) rostral, (B) mid and (C) caudal extent of the NTS. n=3-4/group. *P<.05, **P<.01, ***P< .001, ****P<.0001. Abbreviations: *NTS,* nucleus tractus solitarius.

### Identification of vagal sensory input originating from the stomach and their associated central projections

Five days following the injection of H129-RFP injection into the gastric fundus in sham- and VSG-operated, unipolar H129 labelled cell bodies were identified in the nodose ganglia on both left and right sides with the number of infected cells displaying similar variability between rats in both sham (n=5) and VSG-operated (n=5) groups (Fig.6a). H129-labelled neurons were present throughout the rostro-caudal extent of the NTS. In the most caudal part, labelling was detected at the commissure of the caudal NTS where labelling extended bilaterally throughout the left and right NTS (Fig.6b). In cases where infection was more advanced, labelling extended along the ventrolateral boundary of the NTS into the dorsal motor nucleus of the vagus (DMV-X; Fig.6c) as well as the area postrema. Clusters of infected cells were also found in the locus coeruleus (LC, not shown) and the medial and lateral parts of the PBN (Fig.6d).

**Figure 6.**
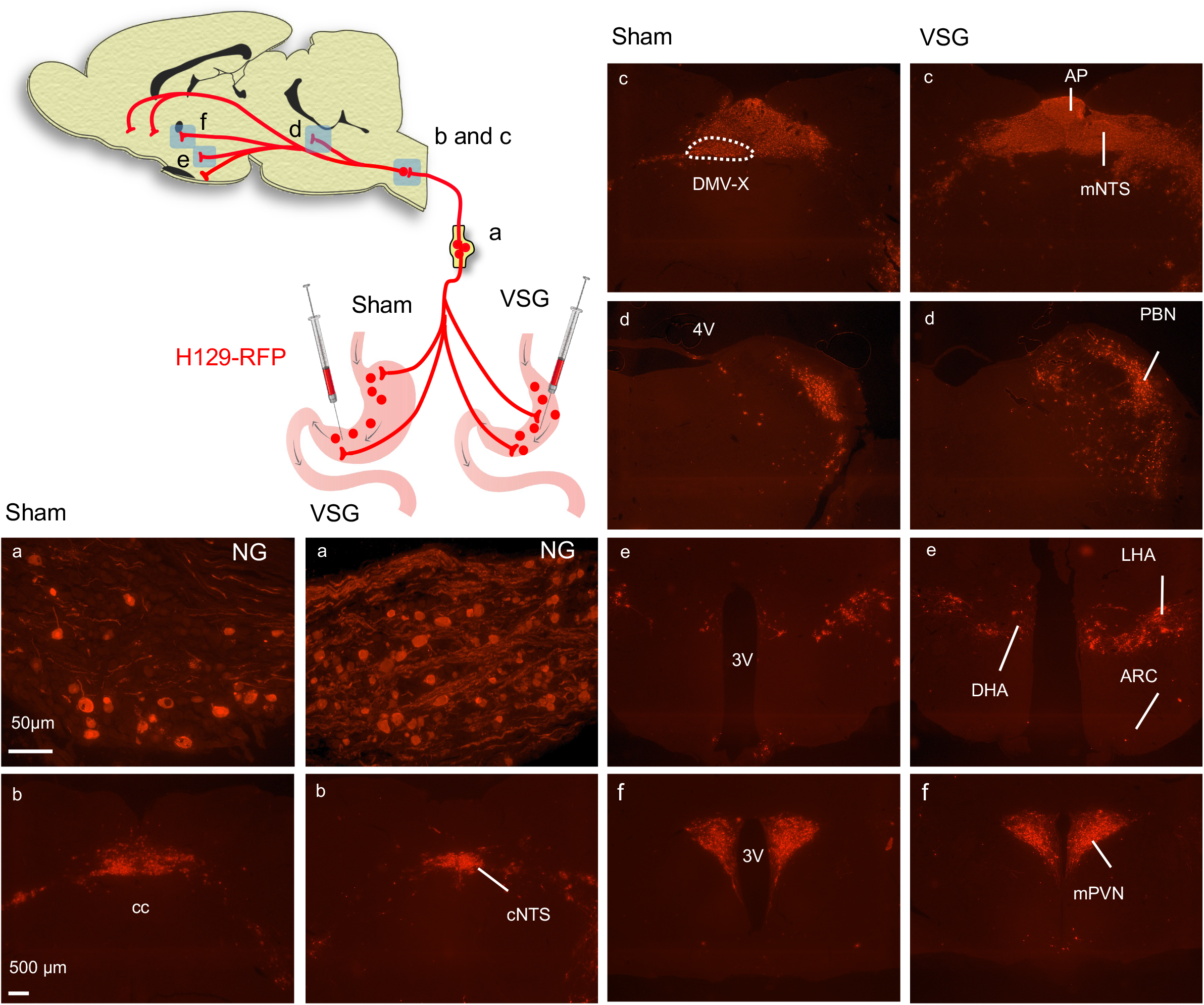
Photomicrographs illustrating the distribution of H129-labelled cells in coronal sections through the caudal to rostral span of the brainstem and forebrain regions of H129-infected sham (column 1,3) and VSG-operated rats (column 2,4) (a) nodose ganglion (NG), (b) commissural NTS, (c) mid NTS, (d) parabrachial nucleus (PBN), (e) dorsal hypothalamic area (DHA), lateral hypothalamic area (LHA) and arcuate nucleus (ARC) and (f) mid paraventricular nucleus of the hypothalamus (mPVN). Scale bar=500 μm. Abbreviations: 3V=3rd ventricle; 4V=4^th^ Ventricle, DMV-X=dorsal motor nucleus of the vagus.

Viral labelling also extended to into forebrain regions including the hypothalamus and POAs. Viral labelling was similarly noted in sham- and VSG-operated rats in the BnST and the medial POA, as well as the central nucleus of the amygdala and the OVLT (data not shown). Moderate H129 viral labelling was present in the dorsomedial hypothalamic nucleus, lateral POA, dorsal hypothalamic and lateral hypothalamic area (Fig.6e) in all animals. In addition, labelling in the parvocellular paraventricular neurons was strongly present throughout the rostro-caudal divisions and this was uniformly bilateral in both sham- and VSG-operated rats (Fig.6f). Following longer survival, there was evidence of viral labelling present in the arcuate nucleus, albeit only a few cells per section in both sham- and VSG-operated rats.

### Identification of synaptic relays between sensory pathways from the stomach or small intestine and efferent outflows to the BAT

#### Relays originating in the stomach

Early PRV-GFP labeling, consistent with premotor or 3^rd^ order projections to the brown fat were found in the LC (not shown), the raphe, RVMM and RVLM of the brainstem, and the PVN, LHA and RCA of the hypothalamus. At longer survival times following injection of PRV-GFP into BAT, an additional order of neurons was labelled including the NTS in the brainstem and the arcuate nucleus in the mediobasal hypothalamus and OVLT in the preoptic region. In the same rats, H129-RFP was detected in overlapping regions that contained PRV-GFP positive neurons with the most predominant labeling in the raphe, RVMM and RVLM as well as the PVN and RCA of the hypothalamus (Fig.7A, Supplementary Table 1). The greatest colocalisation of the two viruses (PRV and H129), consistent with a patent synaptic relay between the stomach and brown fat, was found through the dorsal raphe and raphe pallidus with fewer double labelled neurons in the RCA and limited overlap in the PVN (Fig.7A, Supplementary Table 1).

**Figure 7.**
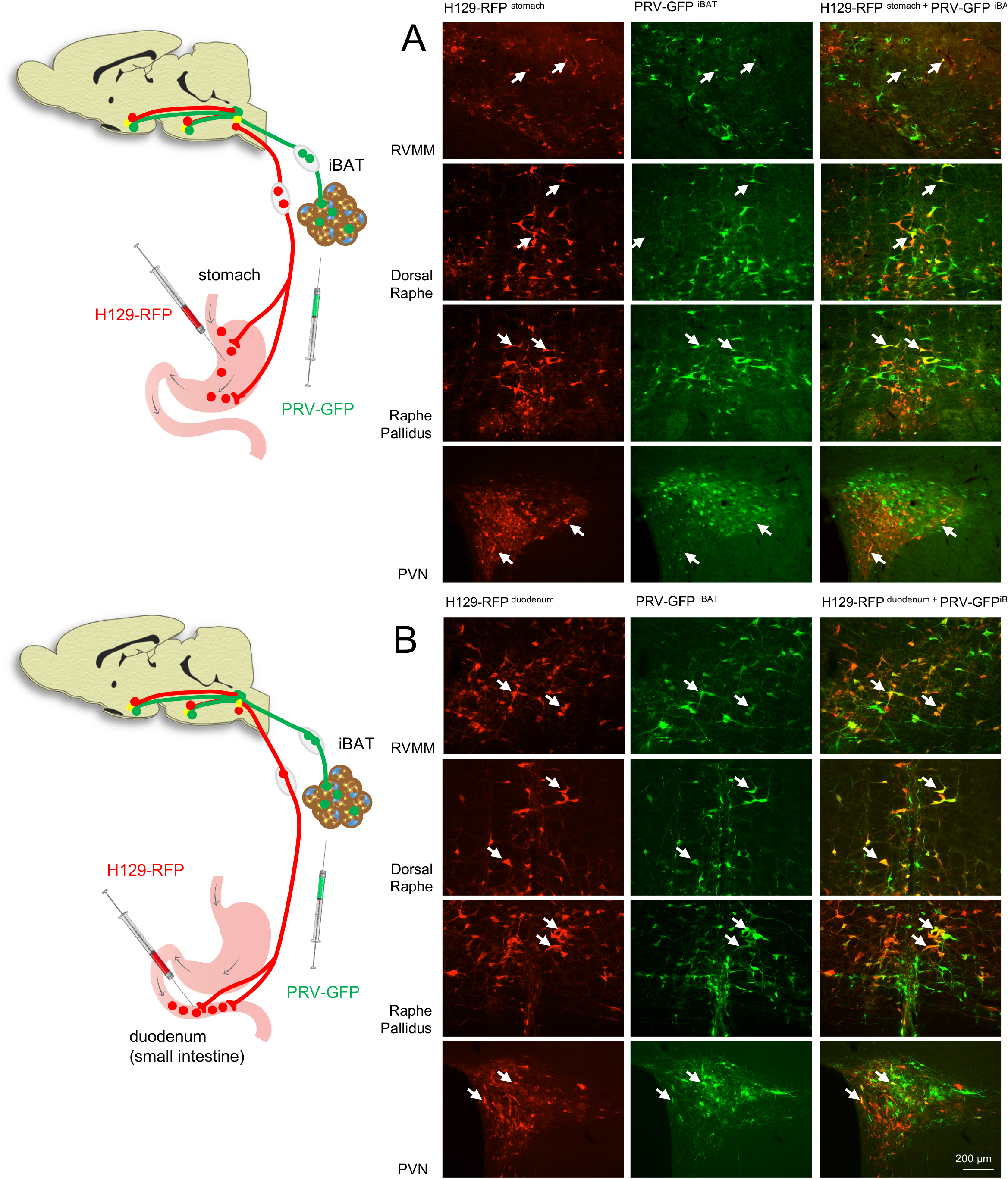
Photomicrographs of H129-RFP and PRV-GFP positive neurons within premotor regions in the brainstem and hypothalamus following injection of (A) H129-RFP into the stomach and PRV-GFP into iBAT and (B) H129-RFP into the small intestine and PRV-GFP into iBAT in the same rat. Co-localization of projections are seen in the rostroventromedial medulla (RVMM), dorsal raphe, raphe pallidus and paraventricular nucleus of the hypothalamus (PVN). Arrows indicate neurons receiving sensory information from the stomach or small intestine and projecting polysynaptically to both iBAT.

#### Relays originating in the duodenum

The hypothalamic and brainstem labeling following PRV injection into the BAT was consistent between the two procedures and similarly, the distribution of sensory neurons following injection of H129-RFP into the duodenum was similar to that observed from the stomach with a (qualitatively) similar density of labeling. Importantly, despite similar labeling patterns from the two injections, the extent of co-existence of viruses within specific regions was far greater, particularly in the RVMM, RVLM, dorsal raphe and raphe pallidus after duodenal injections of H129-RFP (Fig.7B, Supplementary Table 1); consistent with a greater preponderance of synaptic relays derived from the small intestine and directed to the BAT.

## Discussion

These data, derived from a rodent model, further elucidate the mechanism of action of vertical sleeve gastrectomy and its role in mediating weight loss. The findings recapitulate the well-documented effects of VSG surgery on the reduction of food intake, body weight and fat mass^14, 26^. The observation that pair-feeding did not result in the same trajectory in reduction in body weight loss as VSG surgery, specifically over the first 10 days, indicates that reduced food consumption does not solely account for the rapid weight loss following bariatric surgery. This, in conjunction with the observation that sustained weight loss induced by VSG persists despite similar intake between sham and VSG-operated rats over the latter part of the measurement period indicates that increased energy expenditure may contribute to both the rapid weight loss and maintenance of weight loss following VSG. Indeed, VSG was associated with a significant increase in BAT thermogenesis, as demonstrated by an elevation in iBAT temperature and increased UCP1 expression relative to sham controls. The contribution of BAT thermogenesis to VSG-induced weight loss was further substantiated by the chemical or surgical denervation of BAT, after which VSG-induced weight loss was reduced by 50%. In addition, coincident with VSG, there was an elevation in the expression of genes associated with “beiging” of white adipose tissue. Collectively, these data support a role for increased thermogenesis in BAT and possibly in the browning of white fat depots in VSG-mediated weight loss.

While the present study has focussed on VSG, the question arises as to whether there is a common mechanistic basis for the efficacy of related resectional procedures such as RYGB where metabolic outcomes in weight loss and glycemic control are similar. Despite some differences in nutrient processing and gastrointestinal hormone release following RYGB and VSG, restructuring the stomach in each case results in enhanced delivery of nutrients and bile to the jejunum/ileum. The net effect is elevated expression of markers of activity of either, or both, classical BAT or BeAT arising from RYGB and VSG, albeit to different degrees.

A recent review claimed that in both clinical studies and rodent models, there is minimal recruitment of “classical” BAT after RYGB with any post-surgical increase in thermogenic activity restricted to BeAT^18^. This is likely an oversimplification as in some cases where elevated BeAT activity was highlighted, BAT was not examined^27, 28^ and in others where there was no reported “increase” in BAT thermogenesis after RYGB, there was in fact prevention of the reduction in energy expenditure seen in weight matched controls^29^. Moreover, when BAT was examined specifically after RYGB, in both patients with obesity^21^ and in obese mice^30^, activity was elevated as measured by increased uptake of fatty acids into BAT in humans or expression of molecular markers of BAT function in mice.

The recruitment of BAT and BeAT is only slightly more consistent following VSG. Here, similar to data derived from RYGB in rats^29^, total energy expenditure was variably affected after VSG^22, 31^ but even when it was not increased, VSG could prevent the reduction in energy expenditure and BAT-derived UCP1 evident in body weight matched controls and occurred commensurate with an increase in fatty acid utilisation in BAT^31^. In a similar vein, studies in streptozotocin-induced diabetic rats found that VSG caused an adiponectin and SIRT1-mediated browning of inguinal fat and subsequent activation of AMPK to upregulate a raft of thermogenic genes including PGC1α and UCP1^32^. These studies support a positive impact of VSG in both BAT and the browning of WAT following VSG and are coincident with either reduced body weight or an improvement in glucose regulation and reduction of triglycerides. This is consistent with the established role of activated brown and beige fat^33^ but no causal link was established. In this regard, preferential uptake of 18-FDG into inguinal and epidydimal WAT after VSG in rats may help explain the surgery-induced improvement in glucose regulation but, in this case, BAT was not examined^34^. These data are consistent with those derived from clinical studies where elevation of activated BAT^21, 35^ and BeAT^36, 37^ occur after VSG, again contemporaneously with improved glucose regulation^21^.

Considering these data together, there is converging evidence that, firstly, BAT and BeAT are recruited by poorly defined mechanisms following RYGB and VSG and secondly, there is an association between this elevated thermogenic activity and improvements in glucose regulation. Given that BAT and BeAT represent a sink that contributes inordinately to the sequestration of circulating glucose and triglycerides^38^ and the heightened activity of these thermogenic depots after surgery, these tissues may represent, at least in part, the elusive connection that drives the rapid improvement in glucose regulation after RYGB and VSG. Importantly, our data show for the first time a causal link whereby positive metabolic adjustments after surgery can be reversed by the denervation or extirpation of BAT. Moreover, in obese rats, our results demonstrate a VSG-mediated increase in glucose uptake specifically into BAT (and possibly BeAT) and does not alter the uptake into other key metabolic tissues where VSG impressively normalises glucose uptake into the BAT of the obese rat to that of its lean chow-fed controls.

With respect to how changes in the structure and function of the gut following VSG may elicit upregulation of thermogenic tissues, there are a number of possibilities including a blood-borne route for gut-derived factors or hormones including secretin, given its capacity to drive sequestration of glucose by BAT^39^ via direct actions on BAT. Alternatively, the recruitment of vagal afferent neurons (VANs) seems the most plausible based on current evidence. As shown here and elsewhere^14^, VSG in rodent models results in elevation of neural activity (Fos expression) in the first synaptic relay of VANs in the NTS (see also^14^), likely due to the increased intraluminal pressure in the sleeved stomach. In this regard, it has been shown that VANs, possibly characterised by their expression of oxytocin receptor (Oxtr) with endings distributed throughout the stomach and duodenum, are likely candidates to transfer mechanosensitive (pressure sensitive) information in the vagus^40^. However, the relative contribution of mechano- vs chemo-responsive VANs to the efficacy of VSG has remained unclear. Even under very similar experimental conditions of VSG in rats, experiments have described different relative responses to nutrients and stretch. In the present study, a standard volume of high caloric intralipid generates approximately double the number of activated neurons in the NTS than the same volume of a non-caloric (water) load, consistent with an additive impact of both stretch and nutrient modalities on VSG outcomes; however, others have attributed this elevated neural activity solely to an enhanced sensitivity to nutrients^14^.

The dual viral tracing approach employed in the present study highlights, for the first time, a patent stomach / duodenal-brain-BAT neural pathway with relays predominantly through the raphe nuclei in the rostral medulla. This conduit involving VANs innervating the stomach and duodenum might easily subserve the activation of stretch or chemoreceptors, or the recognition of changes in bile acids all of which have been shown to change after VSG and potentially impact the vagus nerve (for review see^41^). In regard to the latter, bile acids are well known to be elevated following RYGB and VSG^42^. Moreover, bile acids robustly stimulate the release of incretins, GLP-1 and GIP, as well as other hormones derived from enteroendocine cells including PYY, via an action on the TGR5^43^. The expression of TGR5 increases in the ileum after VSG with a time course coincident with improved glycaemic control and increased energy expenditure, an effect that is ameliorated in TGR5-KO mice^23^. Of relevance to the proposed gut-brain-BAT reflex, bile acids also increase energy expenditure by their direct actions on brown adipocytes, preventing obesity and insulin resistance^44^. We do not know of any data that would help discriminate between these possible TGR5-mediated pathways underlying the actions of bile acids through either direct incretin effects on insulin production, incretin (GLP-1) actions on vagal afferent neurons, or bile acids actions directly on the BAT. Our data and that of others^14^ showing the activation of VAN relays in the NTS after VSG would suggest that there is at least a contribution to the sensing of VSG-induced perturbations in the gut, and transmission to the brain via the vagus nerve, of modalities that may involve BAs, hormonal, chemo- and mechano-receptor activation. Moreover, the amelioration of the body weight and glycaemic benefit of VSG after BAT denervation is further consistent with the efferent (sympathetic) limb of this proposed neural loop between gut and BAT responsible for the recruitment of BAT and improvements in metabolic outcomes post VSG.

The present study refines the gathering but disparate evidence that points to the pivotal role of BAT in improved metabolic outcomes after VSG. Moreover, it defines a patent neural pathway that may underpin the elusive, short-term improvement in glycemic control after VSG, effectively linking the gut to the BAT via synaptic relays in the brain, particularly via those in the brainstem raphe nuclei. While cognisant of the fact that this is unlikely to represent the only means by which the post-surgical gut communicates with BAT, these data and those of related studies, provide insights that may help mimic the mechanisms of action of surgery-induced metabolic improvements in a non-surgical context.

## Abbreviations

(2-DG): 2 deoxy-D-glucose
(6-OHDA): 6-hydroxydopamine
(BeAT): beige adipose tissue
(BAT): brown adipose tissue
(BnST): bed nucleus stria terminalis
(CNS): central nervous system
(DIO): diet-induced obese/obesity
(DEXA): dual energy x-ray absorptiometry
(FGF-19): fibroblast growth factor-19
(GFP): green fluorescent protein
(GLP-1): glucagon like peptide-1
(GTT): glucose tolerance test
(H129): herpes simplex virus strain 129
(iBAT): interscapular brown adipose tissue
(LC): locus coeruleus
(NTS): nucleus of the solitary tract
(OVLT): Organum vasculosum laminae terminalis
(PYY): peptide YY
(POA): preoptic area
(PRV): pseudorabies virus
(RFP): red fluorescent protein
(RER): respiratory exchange ratio
(RVLM): rostral ventrolateral medulla
(RVMM): rostral ventromedial medulla
(RYGB): roux en Y gastric bypass
(UCP1): uncoupling protein 1
(VANs): vagal afferent neurons
(VSG): vertical sleeve gastrectomy
(WAT): white adipose tissue

## Supplemental Methods

### Pair-feeding

To isolate the impact of VSG directly from that arising secondary to reductions in food intake following surgery, a subset of DIO sham-operated rats were pairfed to *ad libitum* fed rats that underwent VSG. This pairfed group had access to the same amount of food that was consumed by the *ad libitum-fed* VSG group on the preceding day.

### Surgical procedures

Sham and VSG surgery: Rats either underwent sham operation or VSG surgery. Following an overnight fast, rats were anaesthetised using isoflurane (2% in oxygen) (Isorrane, Baxter Healthcare, Australia). Rats were administered the anti-inflammatory meloxicam (2 mg/kg subcutaneously; Boehringer Ingelheim Pty Ltd, North Ryde, NSW, Australia) and long-acting antibiotic oxytetracycline (20 mg/kg; intramuscularly, Animal Health Inc, MO, USA) at the beginning of surgery to control postoperative pain, inflammation and infection. Essentially, the forestomach was completely resected and then sutured with a series of interrupted stitches [4/0 taper polydioxanone (PDS), Dynek, SA, Australia]. The lateral portion of the glandular stomach was then resected and sutured to generate a tubular gastric sleeve, continuous with the oesophagus and duodenum. Overall, the stomach volume was reduced by 70-80 %, a reduction which was verified in post-mortem examination. Sham surgery involved the same abdominal laparotomy incision to expose the stomach, however, no resection was performed.

#### Post-operative care following sham and VSG surgery

Animals were closely monitored during a one-week recovery period following surgery, during which rats were provided with a liquid diet (Ensure, Abbott Nutrition, NSW, Australia) and transitioned back to the high fat diet. During this post-surgical recovery period, animals were administered the anti-inflammatory agent, meloxicam (20 mg/kg subcutaneously; Boehringer Ingelheim Pty Ltd, Australia) daily. In addition, the antibiotic enrofloxacin (Baytril, Bayer Healthcare, Australia) was added to drinking water [1 mL Baytril (25mg/mL)/500 mL of water] throughout the post-surgical recovery period.

#### Surgical and chemical denervation of BAT

For both surgical and chemical denervation procedures iBAT pads were exposed via a dorsal midline incision between the scapulae. Once the borders of both iBAT lobes were identified, a complete excision was made. Chemical denervation using 6-OHDA, a catecholaminergic neurotoxin that has been shown to deplete noradrenergic stores in nerve endings ^45–47^, involved 10 x 1μl injections of a 9 mg/ml 6-OHDA (Sigma Aldrich, North Ryde, NSW, Australia) solution (dissolved in saline) at loci throughout each lobe of iBAT using a specialized syringe (Hamilton Syringe 75 RN 5 μl, model 7634-01, Hamilton Company, NV, USA) connected to a 30G needle (Hamilton model 7803-07, Hamilton Company, NV, USA).

### Changes in body weight, food intake and body composition

Body weight and 24 h food intake were measured daily throughout the experimental period. Total lean, fat and bone mass were assessed using dual energy x-ray absorptiometry (DEXA) prior to surgery and at the end of the experimental period.

### Changes in energy expenditure

#### BAT temperature as a marker of BAT thermogenesis

Rats were surgically fitted with temperature-sensitive probes (TA10TA–F40, DataScience, St. Paul, MN) between the two lobes of interscapular BAT (iBAT). Together with a transmitter device positioned subcutaneously in the flank region, this arrangement enabled measurement of changes in BAT temperature that are reflective of thermogenic activity as well as levels of physical activity in the x–y plane. The signal generated by the probes was detected by receiver plates (RPC–1, DataScience, St. Paul, MN) beneath the rat cages. Following surgical recovery and at a point where VSG operated mice were in their active weight loss phase, both BAT temperature and physical activity were sampled at 5 min intervals continuously for a 48h period.

#### Whole-body energy expenditure using indirect calorimetry

Rats were accommodated to the calorimetry cages (TSE Systems, Bad Homburg, Germany) for at least 24 h (day 1) followed by a 48h measurement period (days 2 and 3). All rats then underwent an overnight fast (18h) and food returned on the following morning (0930h; day 4) and recording continued for the subsequent 3h. Over the entire period, O2 consumption (VO2) and CO2 production (VCO2) were measured every 30 min/cage for 3 min and recorded using specialized software. Respiratory exchange ratio (RER) was calculated as the quotient of VCO2/VO2, with a value of 1.0 representing 100% carbohydrate oxidation, and 0.7 representing 100% fat oxidation.

### Assessment of changes in glucose regulation

#### Oral glucose tolerance testing

Oral glucose tolerance tests were conducted after a six-hour fast (food removed at the onset of the light phase, 0700 h), 30 days after VSG or sham surgery. Oral glucose tolerance was assessed by delivering a 50 % glucose (Glucodin, Rickett and Colman Pharmaceuticals, West Ryde, NSW, Australia) load (1.5 g/kg) via an oral gavage. Blood glucose levels were then assessed by tail prick and determined immediately using a glucometer (Accu-Chek Performa, Roche Diabetes Care, Bella Vista, NSW, Australia) at 0, 15, 30, 60 and 120 min intervals. Blood was also collected from a tail nick (~50 μl) prior to and 15 min following the oral glucose load and plasma insulin was later determined using a commercially available kit (Ultra Sensitive Rat Insulin ELISA Kit (90060), Crystal Chem, IL, USA).

#### Glucose stimulated glucose uptake

To localize glucose uptake in specific tissues we performed an oral GTT (1.5 g/kg) combined with an intraperitoneal injection of [1-14C]-deoxy-d-glucose (40uCi in 200 ul saline). Blood samples were collected before (0) and 2, 5, 10, 20 and 30 minutes after gavage. Tissues were harvested 30 min following oral glucose gavage and snap frozen in liquid nitrogen. Later, radioactivity in blood and tissues were measured using a Beckman Coulter scintillation counter. For blood samples, radioactivity was determined by adding 10 ul plasma to 20 ul ZnSO4 (5 %) (Sigma Aldrich, North Ryde, NSW, Australia), which was topped with 20 ul saturated BaOH (Sigma Aldrich, North Ryde, NSW, Australia) and vortexed immediately. After centrifuging (4000 rpm for 10 min), 15 ul supernatant was combined with 85 ul of water, mixed and topped up with scintillation fluid. Tissues were weighed and homogenised in 1.4 ml ZnSO4 (2.75 %), and 400ul of this homogenate was mixed with 140 ul saturated BaOH. Both homogenates were centrifuged (10 min, 13000rpm and 400 ul of the supernatant were taken and combined with scintillation fluid to assess total and free counts, respectively.

### Assessment of markers of thermogenesis and browning in BAT and BeAT

#### Changes in protein expression using Western Blot analysis

Interscapular BAT pads were removed post-mortem and snap-frozen in liquid nitrogen. Protein from BAT was extracted as previously described^48^. For each animal, 40 μg of the extracted protein was loaded onto a Bio-Rad Mini-PROTEAN TGX Precast Gel (Bio-Rad Laboratories, Hercules, CA) and transferred onto an Immuno-Blot nitrocellulose membrane (Bio-Rad Laboratories, Hercules, CA). The membrane was blocked with skim milk powder in Tris-buffered saline/1% Tween 20 (Sigma, St Louis, MO) for one hour and then incubated overnight at 4°C in a primary antibody raised against uncoupling protein 1 (UCP1, Santa Cruz Biotechnology, Santa Cruz, CA; diluted 1:1000) in 5 % bovine serum albumin in Tris-buffered saline/Tween 20 solution. Following secondary antibody incubation with donkey anti-Goat IgG HRP (Antibodies Australia, VIC, Australia; diluted 1:4000), protein expression signals were visualized by chemiluminescence using the Pierce ECL Western Blotting Substrate (Thermo Scientific, Rockford, IL) on the Bio-Rad ChemiDoc system (Bio-Rad Laboratories, Hercules, CA). Relative densities of protein bands were assessed using Image Lab software (National Institutes of Health, Bethesda, MD).

#### Changes in protein expression using quantitative RT-PCR

Total RNA was extracted from inguinal white adipose tissue (iWAT) with Trizol reagent (Thermo Fisher Scientific, Waltham, MA, USA) and its quantity determined using a NanoDrop 3300 (Thermo Fisher Scientific). mRNA was reverse transcribed into cDNA (iScript Reverse Transcription Supermix for Quantitative RT-PCR; Bio-Rad Laboratories, Hercules, CA, USA), and gene expression was determined by real-time qPCR using the TaqMan Fast Advanced Master Mix (Product: 4444557, Applied Biosystems by Thermo Fisher Scientific, MA, USA) and TaqMan Gene Expression Assays (Applied Biosystems by Thermo Fisher Scientific, MA) for UCP1 (Rn00562126_m1), Dio2 (Rn00581867_m1), Ppargc (Rn00580241_m1), Cpt1b (Rn00682395_m1), Cox8b (Rn00562884_m1), Cited1 (Rn00821880_g1), Tbx1 (Rn01405401_m1), Tnfrsf9 (Rn01519016_m1), Tmem26 (Rn01428021_m1), Cidea (Rn04181355_m1), Eno1 (Rn008200594_g1), Pkm (Rn00583975_m1) and 45S (Rn.03928990_ g1). Relative gene expression was calculated using the 2^−ΔΔCt^ method.

### Neural activation following acute infusion of either nutrient or non-nutrient loads into the VSG stomach

#### Preparation of brain sections for immunohistochemistry

Brains were removed, fixed and cut in a coronal plane at 35 μm using a cryostat (Leica CM3050 Research Cryostat, Leica Biosystems, Australia). In order to identify activated neurons, sections were exposed to Fos antisera (Santa Cruz Biotechnology, Santa Cruz, CA; rabbit anti c-Fos, sc-52, 1:1000) that were localized with an immunoperoxidase method resulting in a brown reaction product in reactive nuclei.

#### Cell counts

The extent of labelling of Fos-positive neurons was quantified in specific brain regions in the brainstem. Blind counts of labelled neurons were made through the rostrocaudal extent of the nucleus tractus solitarius (NTS), and divided into rostral (12.00 to 13.00 mm from bregma), mid (13.00 to 14.00 from bregma) and caudal (14.00 to 14.60 mm from bregma) portions.

### Injection procedures for neuroanatomical tracing

#### Anterograde tracing using herpes simplex virus-1

In order to conduct neuroanatomical tracing of VANs and their neural connections, a recombinant of the H129 that expresses red fluorescent protein (RFP; H129-RFP) was used. The viral infection procedure was performed immediately after performing sham or VSG surgery. Using a 10μl Hamilton (model 7635-01, Hamilton Company, NV, USA) syringe with a sharpened beveled tip, 20 x 0.5 μl injections were made within the glandular portion of the stomach (or duodenum, see below). The injections were made on both the dorsal (x10) and ventral (x10) surfaces of the lesser curvature of the stomach of both sham (n=5) and VSG (n=5) operated animals. In the case of the sham-operated rats, care was taken to ensure the virus was directed to the corresponding region of the stomach that formed the “sleeve” in VSG rats. Following each injection, the syringe was held in place for at least 30 seconds to allow for the diffusion of the virus from the needle tip into the injection site, and to minimise efflux and contamination of adjacent tissues. Following the last injection, the stomach was rinsed with saline and swabbed dry to remove any residual virus, after which the stomach was returned to the abdominal cavity and muscle and skin incisions closed. Animals were allowed to survive for up to five days post-inoculation to ensure early arrival of the virus at the region at which point they were perfused transcardially with saline and 4% paraformaldehyde and nodose ganglia and brains collected from all animals.

#### Dual viral approach – combination of H129 and PRV injections

In order to identify synaptic relays between sensory pathways from the stomach or small intestine and efferent outflows to the BAT, a combination viral approach was adopted that involved the injection of H129-RFP into the stomach or small intestine and PRV-GFP into the iBAT in the same animal. Injections into the stomach or duodenum were first performed (as above) to identify sensory pathways in the discrete brain regions. It was necessary to stagger the injection of the iBAT by one day to ensure simultaneous arrival of both viruses within spinally-projecting (premotor, 3rd order) and higher (4th order) regions. Therefore, one day following injection of H129-RFP in the stomach or duodenum, the iBAT was exposed through a midline dorsal incision between the two scapulae. Using a Hamilton syringe (model 7634-01) with a sharpened beveled tip, 8x 0.5μl injections (4 × 0.5 μl injections per lobe) of PRV-GFP were made into the iBAT. Following the final injection in each series, the syringe was withdrawn, and the injection site was padded dry to remove any residual virus. Four days following viral inoculation of the iBAT, rats were perfused with aldehydes and brains collected. In order to better visualise the signals of the fluorescent signals associated with the H-129 virus or PRV, free floating sections of the brainstem and forebrain as well as slide mounted nodose ganglia were subjected to standard immunohistochemical protocols using primary antibodies for 24 h at room temperature for GFP (chicken anti-EGPF, 1:2000, Abcam) and RFP (Living Colors rabbit anti-DsRed, 1:1000, Takara Bio, CA, USA).

#### Tissue analysis, imaging and figure preparation

Mounted nodose ganglia and brains were analysed using fluorescent microscopy (Axio Imager Z1, Zeiss, Germany). Images were captured and processed using a digital camera (AxioCam MRm v4.5, Zeiss, Germany) and digital image processing software (AxioVision v4.8.2, Zeiss, Germany). Comparisons of the distribution of H129-positive neurons within the nodose ganglia and brains following either 1) H129-RFP injection into the stomach or 2) H129-RFP injection into the stomach or duodenum and PRV-GFP into the brown fat.

---

**Supplemental Table 1.**
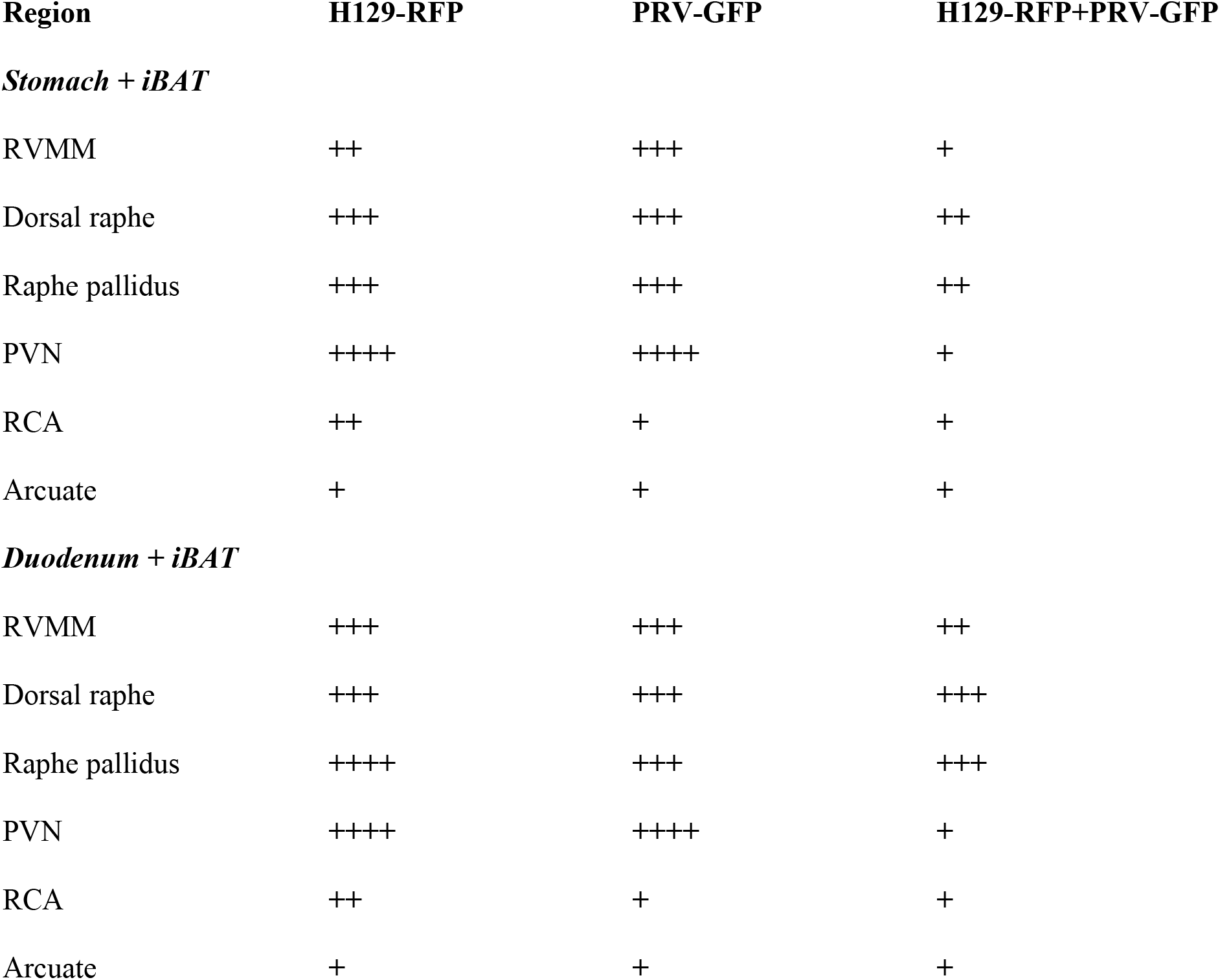
Distribution of H129-RFP following injection into the stomach or small intestine (red) and PRV-GFP following injection into the iBAT in the same rat.

| Region | H129-RFP | PRV-GFP | H129-RFP+PRV-GFP |
| --- | --- | --- | --- |
| *Stomach + iBAT* |  |  |  |
| RVMM | ++ | +++ | + |
| Dorsal raphe | +++ | +++ | ++ |
| Raphe pallidus | +++ | +++ | ++ |
| PVN | ++++ | ++++ | + |
| RCA | ++ | + | + |
| Arcuate | + | + | + |
| *Duodenum + iBAT* |  |  |  |
| RVMM | +++ | +++ | ++ |
| Dorsal raphe | +++ | +++ | +++ |
| Raphe pallidus | ++++ | +++ | +++ |
| PVN | ++++ | ++++ | + |
| RCA | ++ | + | + |
| Arcuate | + | + | + |

The number of H129-RFP and PRV-GFP -positive neurons were qualitatively estimated in the brains of rats. Extent of viral labelling is presented as follows: ++++ (very high: more than 200 -RFP/GFP positive neurons per brain region); +++ (high: between 100 and 200 -RFP/GFP positive neurons per brain region); ++ (moderate: between 20 and 100 -RFP/GFP positive neurons per brain region); + (low: less than 20 -RFP/GFP positive neurons per brain region).

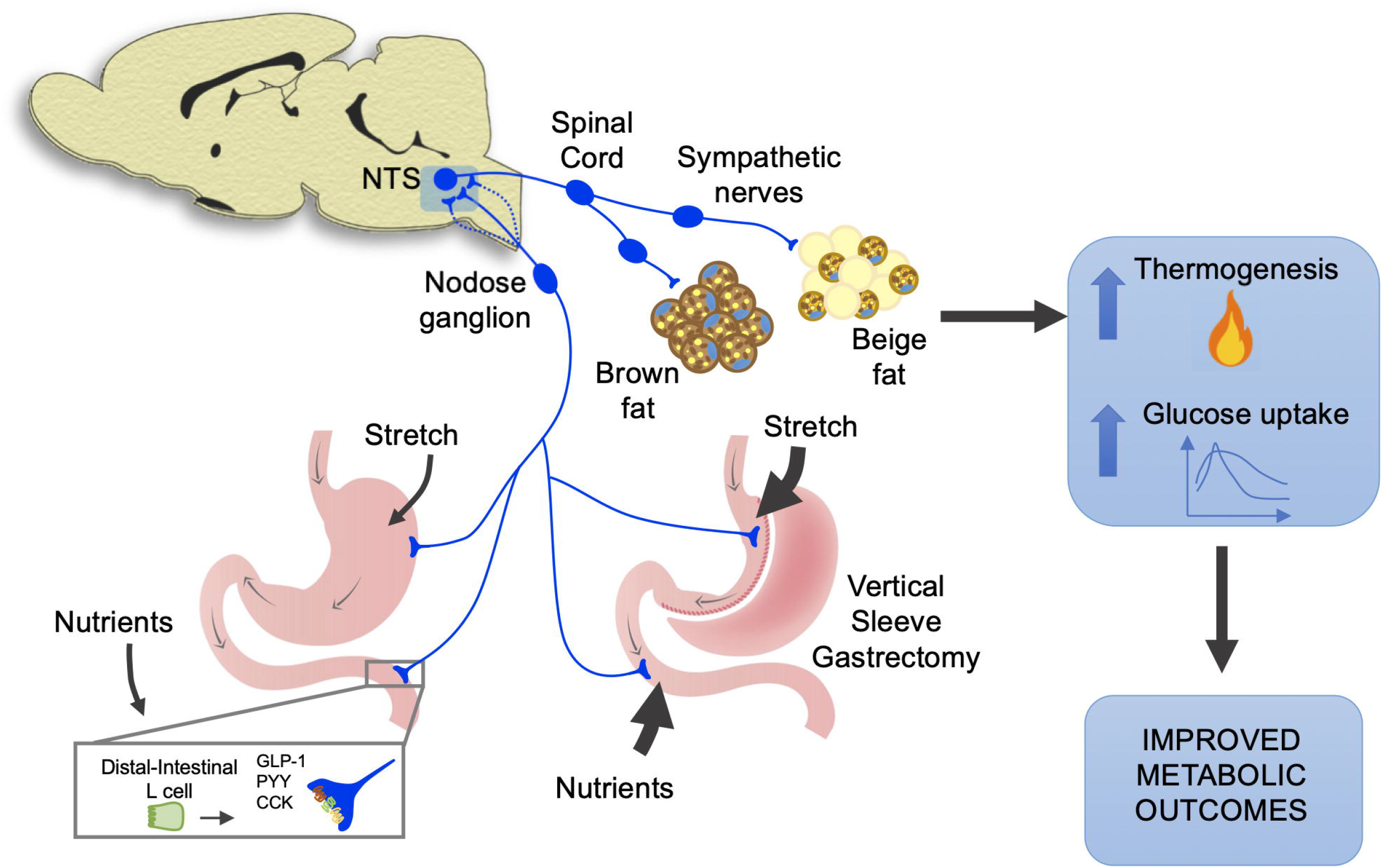

